# Multi-omics analysis reveals the mechanism of BnC07MYB3a is involved in seed coat color in *Brassica napus* L

**DOI:** 10.1101/2024.09.30.615900

**Authors:** Ran Hu, Mengzhen Zhang, Shulin Shen, Haijing Liu, Lei Gao, Mengjiao Tian, Yiwei Liu, Huafang Wan, Huiyan Zhao, Nengwen Yin, Hai Du, Liezhao Liu, Kun Lu, Jiana Li, Cunmin Qu

## Abstract

In rapeseed (*Brassica napus*), yellow-seeded varieties accumulate less flavonoid pigments (anthocyanins/proanthocyanidins) in their seed coats compared with black-seeded varieties. The yellow-seeded trait is associated with greatly improved seed oil yield, quality, and commercial value. Many R2R3 MYB activators have been characterized in rapeseed, but how MYB-type repressors affect pigment biosynthesis is not yet fully understood. In this study, we performed transcriptome sequencing and metabolomic analysis of *B. napus* varieties with extreme differences in seed coat color, combined with weighted gene co-expression network analysis. This analysis identified an R2R3-MYB-type transcription factor, BnC07MYB3a (BnaC07G0178800ZS), as a candidate regulator of the yellow-seeded trait in *B. napus*. Overexpressing *BnC07MYB3a* in *Arabidopsis thaliana* and *B. napus* downregulated the expression of flavonoid biosynthetic genes, resulting in significantly lower anthocyanin and proanthocyanidin accumulation than in the wild-type and a lighter seed coat color in transgenic plants. BnC07MYB3a directly binds to the promoter of the *TRANSPARENT TESTA* (*TT*) gene *BnTT6* and represses its expression. BnC07MYB3a also physically interacts with BnA06bHLH92a and the WD40 transcription factor TRANSPARENT TESTA GLABRA1 (BnTTG1), suggesting that they might form a previously unidentified MYB–bHLH–WD40 transcription factor complex. Our results reveal the molecular mechanism and regulatory network of BnC07MYB3a in determining seed coat color in *B. napus* and provide a genetic resource for breeding yellow-seeded cultivars of *B. napus*.

## Introduction

Rapeseed (*Brassica napus* L.) is an important oilseed crop, providing vegetable oil for the human diet and oil for biodiesel production and serving as a source of protein-rich feed for livestock (Hu et al. 2017; Kimber et al. 1995). The yellow-seeded trait in *B. napus* is commercially valuable as it is associated with more transparent oil, lower fiber content, a more desirable aroma, and higher nutritional content than the black-seeded trait (Tang et al. 1997; Rahman et al. 2001). Given its commercial value, exploring the mechanism underlying the yellow-seeded trait has been a major focus of research over the past three decades. However, natural yellow-seeded rapeseed does not exist: yellow-seeded materials are mainly obtained through distant hybridization or physical or chemical mutagenesis. This seed color trait is genetically unstable and susceptible to environmental factors (Zhi-wen et al. 2005; Deynze et al. 1993; Rashid et al. 1994).

Seed coat color is associated with the accumulation of metabolites of the flavonoid biosynthetic pathway (Qu et al. 2013). Flavonoids are a large group of phenylpropanoid compounds that includes flavones, flavonols, anthocyanins, and proanthocyanins (Winkel-Shirley et al. 2001). In Arabidopsis (*Arabidopsis thaliana*), more than 20 genes related to this pathway have been isolated (Albert et al. 2014), including eight structural genes encoding catalases (*TT3*–*TT7*, *FLS1*, *TT18*, and *BAN*); the early biosynthesis genes (EBGs) *CHALCONE SYNTHASE* (*CHS/TT4*), *CHALCONE ISOMERASE* (*CHI/TT5)*, *FLAVANONE 3-HYDROXYLASE* (*F3H/TT6*), *FLAVANONE 3′-HYDROXYLASE* (*F3′H/TT7*), and *FLAVONOL SYNTHASE* (*FLS*), which mainly regulate the biosynthesis of flavonoid precursors; the late-biosynthesis genes (LBGs) *DIHYDROFLAVONOL-4-REDUCTASE*(*DFR/TT3*), *LEUCOANTHOCYANIDIN DIOXYGENASE* (*LDOX*/*TT18*), and *BANYULS* (*BAN*), which are responsible for anthocyanin biosynthesis (Chiu et al. 2010; Albert et al. 1997; Wan *et al*.; 2002; Routaboul et al. 2006); six regulators of the TRANSPARENT TESTA (TT) and TRANSPARENT TESTA GLABRA (TTG) families (*TT1*, *TT2*, *TT8*, *TTG1*, *TTG2*, and *TT16*) that act on structural genes to regulate the flavonoid pathway, either individually or in complexes (Baudry et al. 2006; Nesi et al. 2000; Nesi et al. 2001; Routaboul *et al*.; 2006); and transporter genes (*TT9*, *TT12*, and *TT19*, among others) encoding proteins involved in proanthocyanidin accumulation, transport, and oxidation (Baxter et al. 2005; Debeaujon et al. 2001). Among these, TT2 (R2R3-MYB), TT8 (basic helix-loop-helix, bHLH), and WD40 transcription factors (WDRs) form ternary MYB–bHLH–WDR (MBW) complexes, which influence the flavonoid biosynthesis pathway by regulating the expression of EBGs and LBGs, leading to changes in seed coat color (Nesi et al. 2001; Xu et al. 2015).

Most of these genes are also associated with the flavonoid biosynthesis pathway in various *Brassica* species: *TTG1* in *Brassica rapa* (Zhang et al. 2009) and *B. napus* (Cheng et al. 2024); *TT8* in *B. rapa* (Li et al. 2012), *Brassica juncea* (Padmaja et al. 2014), and *B. napus* (Zhai et al. 2020); *TT1* in *B. napus* (Lian et al. 2017; Li et al. 2024); *TT2* in *B. napus* (Xie et al. 2020; Li et al. 2024); *TT7*, *TT18*, *TT10*, and *TT12* in *B. napus* (Li et al. 2024); and *MYB47* in *B. napus* (Qu et al. 2023). The expression patterns of these genes differ between yellow- and black-seeded *Brassica* lines (Saigo et al. 2020; Qu et al. 2013; Zhang et al. 2020a; Zhang et al. 2020c; Qu et al. 2023).

R2R3 MYB transcription factors are major players in flavonoid biosynthesis during fruit coloration in many species. Some R2R3 MYB transcription factors also promote the accumulation of anthocyanins and proanthocyanidins, such as AtMYB75, AtMYB90, AtMYB113, and AtMYB114 in Arabidopsis (Onkokesung et al. 2014; Gonzalez et al. 2008); MdMYB1, MdMYB10, and MdMYB110a in apple (*Malus domestica*); and PsMYB134 and PsMYB115 in *Populus* spp. (Mellway et al. 2009; James et al. 2017). The roles of these transcription factors in regulating flavonoid biosynthesis depend on bHLH or WD40 proteins (Wang et al. 2020). In Arabidopsis, the AtMYB114-AtTT8 and TT2-TT8-TTG1 complexes regulate the expression of LBGs (*DFR*, *ANS*, and *BAN*, among others) (Li et al. 2016; Baudry et al. 2004). VvMYB15 promotes anthocyanin accumulation by interacting with VvWRKY40 in grape (*Vitis vinifera*) berries (Li et al. 2021), and also interacts with miRNAs, lncRNAs, COP1, and other molecules (Cao et al. 2016). The SUMO E3 ligase SIZ1 mediates the sumoylation of MYB75, which in turn affects anthocyanin accumulation (Zheng et al. 2020). The E3 ubiquitin ligase COP1/SPA interacts with PAP1 and PAP2, which are required for anthocyanin production.

In addition to MYB activators, several MYB repressors affecting anthocyanin and proanthocyanidin accumulation have been identified in a wide range of plant species. These include AtMYB3 (Kim et al. 2022), AtMYB4 (Banerjee et al. 2024), and AtCPC (Zhu et al. 2009) in Arabidopsis, SlMYB7 in tomato (*Solanum lycopersicum*) (Zhang et al. 2023), MdMYB16 in apple (Xu et al. 2017), VvMYBC2-L1 in grapevine (Cavallini et al. 2015), FhMYBx in *Freesia hybrida* (Li et al. 2020), and BnCPC in *B. napus* (Xie et al. 2022). MYB repressors are classified into three types based on the repression motifs in their C-terminal domains: R2-R3-MYBs, with EAR repression motifs; R3-MYBs, with the unique repression motif “TLLLFR”; and CPC-type R3-MYBs, which lack a terminal repression motif. These repressors inhibit MBW complex formation by interacting with bHLH cofactors, resulting in passive repression (Albert et al. 2014; Zhou et al. 2019; Ding et al. 2020). To date, numerous QTLs (Fu et al. 2007; Zhang et al. 2011) and candidate genes (Qu et al. 2013; Lian et al. 2017; Zhai et al. 2020) involved in seed color have been identified in *B. napus*. However, few MYB-type repressors involved in regulating pigment accumulation in *B. napus* have been characterized.

We previously reported that epicatechin and its derivatives play important roles in the formation of seed coat pigments (Qu et al. 2013; Qu et al. 2023). In the current study, by performing weighted gene co-expression network analysis (WGCNA) using these six metabolites as traits, we identified the MYB-like transcription factor BnC07MYB3a, which negatively regulates pigment accumulation. Functional and expression analyses indicated that BnC07MYB3a is a trans-repressor localized to the nucleus that represses the expression of flavonoid biosynthetic pathway genes. This leads to an overall decrease in proanthocyanidin and anthocyanin levels, resulting in a lighter seed coat color. We also determined that BnC07MYB3a directly represses *BnTT6* promoter activity. In addition, BnC07MYB3a, BnA06bHLH92a, and BnTTG1 may form a previously uncharacterized MBW functional complex. Our findings shed light on the transcriptional regulation of the flavonoid biosynthesis pathway in *Brassica* plants.

## Results

### UPLC-HESI-MS/MS and RNA-Seq-based WGCNA identify candidate genes for differential metabolites

Epicatechin and its derivative proanthocyanidins are the main compounds responsible for the formation of seed coat pigments (Qu et al. 2023; Qu et al. 2013). Here, we used ultra-high-pressure liquid chromatography with heated electrospray ionization tandem mass spectrometry (UPLC-HESI-MS/MS) to examine the accumulation of epicatechin and its derivatives (proanthocyanidins; procyanidin B2 ([DP2]-1) and [DP2]-2, procyanidin C ([DP3]-1) and [DP3]-2, [DP4]) in six *Brassica napus* accessions during seed development. The levels of these metabolites varied greatly between yellow- and black-seeded rapeseed accessions starting at 20 days after flowering (DAF) (Fig. 1A). To explore genes that regulate the biosynthesis of these metabolites, we performed transcriptome sequencing of samples at 20, 30, and 40 DAF. We identified 4,540 genes that were differentially expressed in yellow- vs. black-seeded *B. napus* at three stages of development (Fig. 1B). The differentially expressed genes (DEGs) were significantly enriched in the flavonoid biosynthesis pathway, ubiquinone and other terpenoid-quinone biosynthesis, zeatin biosynthesis, and glucosinolate biosynthesis, as revealed by KEGG enrichment analysis (Fig. 1C). These results suggest that there are dramatic differences in the flavonoid pathway during seed development in yellow- vs. black-seeded *B. napus*.

**Figure 1.**
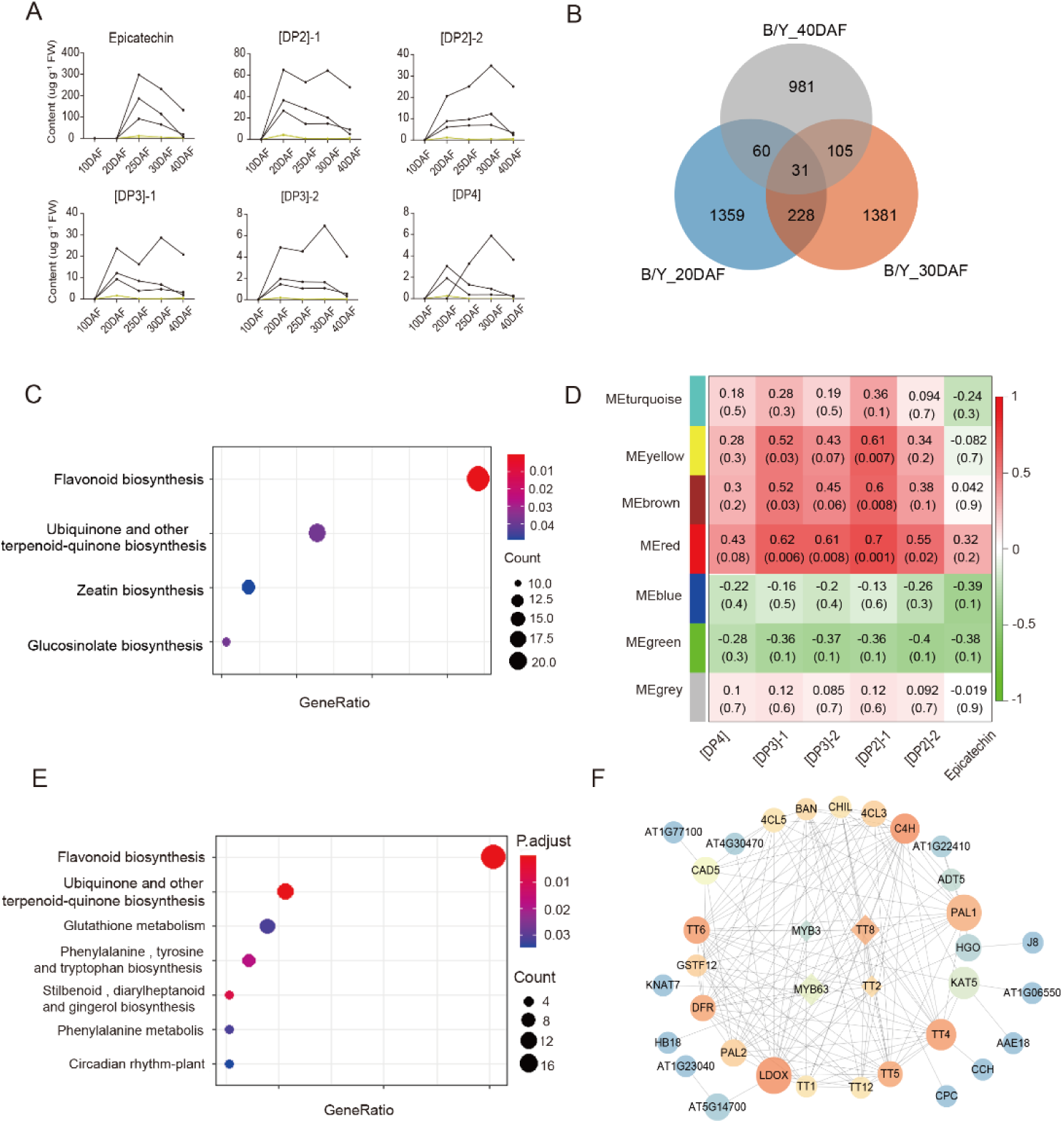
WGCNA identifies BnC07MYB3a, which is associated with differential flavonoid metabolite contents. A. Accumulation patterns of epicatechin and its proanthocyanidin derivatives in seeds at different developmental stages. The black and yellow lines indicate black-seeded and yellow-seeded accessions, respectively. DAF, days after flowering. B. Number of differentially expressed genes (DEGs) in yellow- and black-seeded *B. napus* seeds at different developmental stages. C. KEGG enrichment analysis of all DEGs. D. Heatmap of the relationships of the modules with the levels of epicatechin and proanthocyanidin derivatives. Each cell contains the corresponding correlation and *p-*value in parentheses. E. KEGG enrichment analysis of red module genes. F. Interaction network of red module genes.

Subsequently, we performed WGCNA to identify important regulatory genes. After filtering out DEGs with low expression (average FPKM<0.5), the 1942 remaining DEGs were divided into seven gene modules (Supplemental Fig. S1). Among these, the 163 genes in the red module were significantly associated with the contents of epicatechin and its derivatives (proanthocyanidins; Fig. 1D). Enrichment analysis revealed that these genes were significantly associated with the flavonoid biosynthesis pathway (Fig. 1E). These results suggest that genes in the red module are involved in the flavonoid biosynthesis pathway in rapeseed.

To better understand the roles of genes belonging to the red module in the flavonoid biosynthesis pathway, we constructed protein–protein interaction (PPI) networks to predict the interactions of the proteins encoded by these genes. We identified an interaction network involving multiple flavonoid pathway genes, such as *BnTT8* and *BnTT2* (Li et al. 2024) (Fig. 1F). In addition, we discovered the presence of the MYB transcription factor MYB3, which plays an important role in anthocyanin biosynthesis in Arabidopsis (Kim et al. 2022). However, its function in rapeseed has not been previously reported.

### BnC07MYB3a localizes to the nucleus and is a candidate regulator of the yellow-seeded trait in *B. napus*

The 680 MYB proteins in *B. napus* are divided into four families: 1R-MYB (consisting of one or two separated repeats), 2R-MYB (R2R3-MYB, consisting of two adjacent repeats), 3R-MYB (consisting of three adjacent repeats), and 4R-MYB (consisting of four adjacent repeats). Sequence analysis revealed that BnC07MYB3a contains N-terminal R2 and R3 MYB DNA-binding domains with the conserved motif [D/E]Lx_2_[R/K]x_3_Lx_6_Lx_3_R, for interactions with bHLH proteins. The EAR repression motif (LNL[D/E]L) was also present in BnC07MYB3a, a feature that is consistent with other MYB-type repressors that have been identified (Supplemental Fig. S2A). Phylogenetic analysis grouped BnC07MYB3a with the anthocyanin/proanthocyanidin repressor AtMYB3 (Supplemental Fig. S2B).

The full-length cDNA of *BnC07MYB3a* (BnaC07G0178800ZS) spans 819 bp, encoding a 272 amino acid protein with a predicted molecular weight of 31.127 kDa and a calculated isoelectric point of 8.49 (Supplemental Table S1). Higher levels of *BnC07MYB3a* transcripts were detected in the seed coats of developing seeds of the yellow-seeded accession than in those of the black-seeded accession at 10–20 DAF (Fig. 2A).

**Figure 2.**
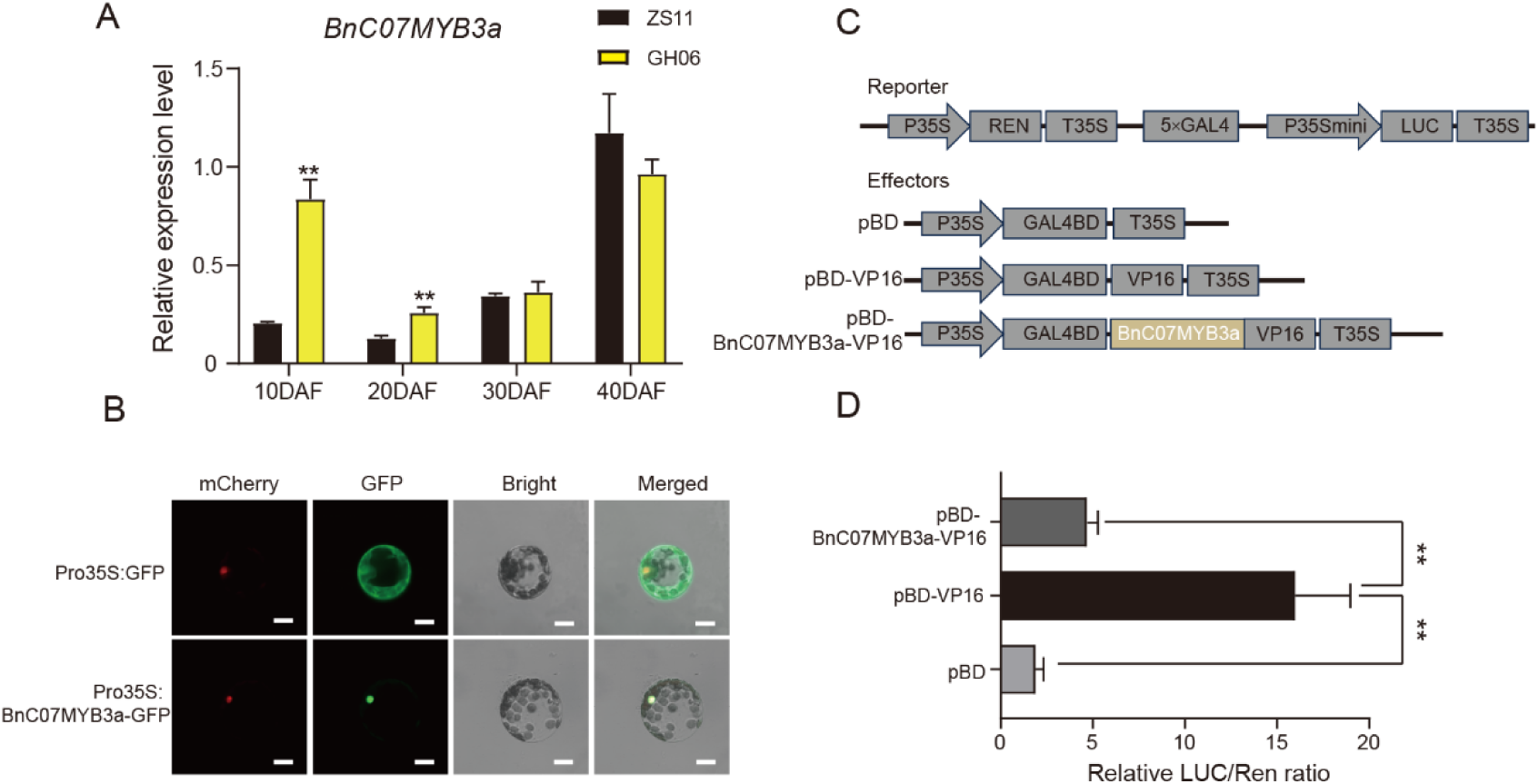
Characterization of BnC07MYB3a. A. Expression patterns of *BnC07MYB3a* in the seed coats of yellow- and black-seeded rapeseed, as revealed by qRT-PCR. Values are mean ± SD of three technical replicates (**, *p* < 0.01). DAF, days after flowering. B. BnC07MYB3a localizes to the nucleus in Arabidopsis protoplasts. BnC07MYB3a-GFP or GFP was transiently expressed in Arabidopsis protoplasts. Bars, 50 μm. C. Diagram of the double-reporter and effector plasmids used in the dual-luciferase assay. D. Transcriptional repression activity of BnC07MYB3a in Arabidopsis protoplasts. Values are mean ± SD of three technical replicates (**, *p* < 0.01). *P* values were calculated using multiple *t* tests without adjustments.

To examine the subcellular localization of BnC07MYB3a, we expressed a BnC07MYB3a-GFP fusion protein under the control of the 35S promoter in Arabidopsis protoplasts. The control (expressing GFP) produced fluorescent signals throughout the protoplasts, whereas the Pro35S:BnC07MYB3a-GFP fusion protein was detected exclusively in the nucleus (Fig. 2B), a finding consistent with its putative role as a transcription factor. To investigate whether BnC07MYB3a is a transcriptional activator or repressor, we fused the full-length cDNA of *BnC07MYB3a* with the VP16 transcriptional activation domain as an effector and expressed this construct in Arabidopsis protoplasts; the VP16 transcriptional activation domain was used as a positive control (Fig. 2C). Protoplasts carrying pBD-BnC07MYB3a-VP16 showed lower expression of the LUC reporter than those carrying pBD-VP16 (Fig. 2D), suggesting that BnC07MYB3a functions as a transcriptional repressor.

### BnC07MYB3a negatively regulates anthocyanin and proanthocyanidin accumulation in Arabidopsis

To identify the biological function of BnC07MYB3a, we constructed an overexpression vector harboring *BnC07MYB3a* driven by the 35S promoter and used it to transform wild-type *A. thaliana* by *A. tumefaciens*-mediated transformation. qRT-PCR revealed highly significant upregulation of *BnC07MYB3a* expression in two T3 lines (OE-*3a#3* and OE-*3a#10*) (Fig. 3A). The seed coat color of the transgenic Arabidopsis plants was lighter than that of the wild-type (Fig. 3B).

**Figure 3.**
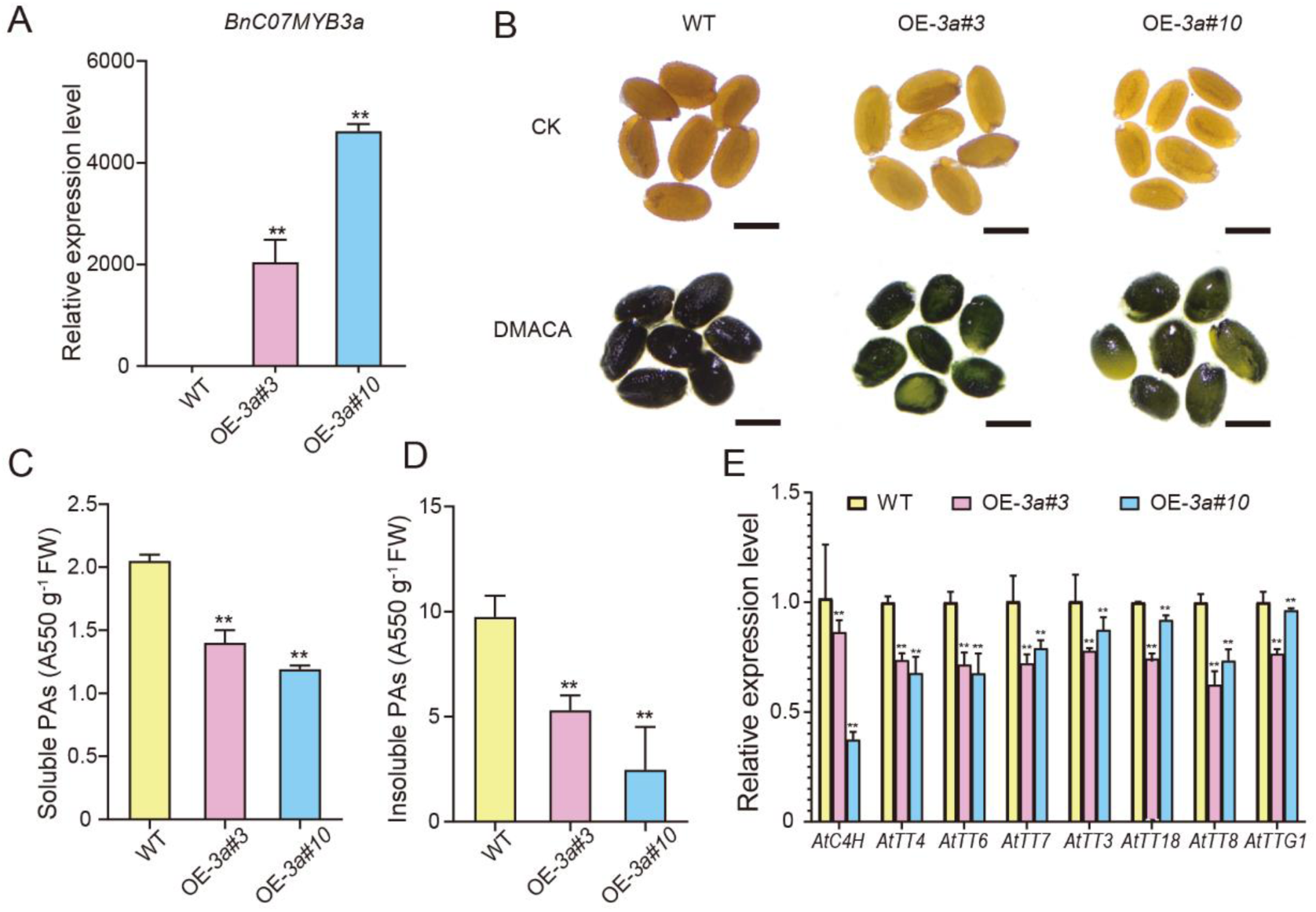
Overexpressing *BnC07MYB3a* inhibits anthocyanin and proanthocyanidin biosynthesis in transgenic Arabidopsis. A. Relative expression levels of *BnC07MYB3a* in *A. thaliana* OE-3a lines. The *A. thaliana ACTIN* gene was used as an internal control. B. DMACA staining of OE-3a and wild-type control seeds. WT, wild type; CK, control; DMACA, 4-dimethylaminocinnamaldehyde. Bars, 1 μm. C. Soluble proanthocyanidin (PA) levels in the WT and two transgenic lines. D. Insoluble PA levels in the WT and two transgenic lines. E. Relative expression levels of flavonoid biosynthesis genes in the WT and in OE-3a Arabidopsis lines, as detected by qRT-PCR. Values are mean ± SD of three biological replicates (**, *p* < 0.01). *P* values were calculated using multiple *t* tests without adjustments.

Proanthocyanidins, which are metabolites that affect seed coat color (Xu et al. 2014; Qi et al. 2011), react with DMACA to produce blue precipitates (Abeynayake et al. 2011; Ferreira et al. 2017). Following DMACA staining, the OE-*3a* lines were a lighter blue color than the wild type (Fig. 3B). Correspondingly, quantitative chemical assays revealed a significant reduction in PA (soluble and insoluble) contents in the OE-*3a* lines compared with the wild type (Fig. 3, C and D).

To examine the effect of BnC07MYB3a on flavonoid biosynthesis pathway genes, we performed qRT-PCR to examine the expression levels of structural and regulatory genes in the OE-*3a* lines. Consistent with the lower proanthocyanidins contents in the OE-*3a* lines, flavonoid biosynthesis genes, including *AtC4H*, *AtTT4*, *AtTT6*, *AtTT7*, *AtTT3*, *AtTT18*, *AtTT8*, and *AtTTG1*, were downregulated in the OE-*3a* lines compared with the control (Fig. 3E). These results indicate that BnC07MYB3a negatively regulates flavonoid biosynthesis.

### BnC07MYB3a negatively regulates anthocyanin and PA accumulation in *B. napus*

The light to dark brown color of mature seeds in *Brassica* species is caused by the oxidation of PAs during seed desiccation, leading to the accumulation of dark brown compounds in the seed coat. To investigate whether BnMYB3a affects PA accumulation in *B. napus*, we obtained transgenic rapeseed overexpressing *BnC07MYB3a* and confirmed the presence of the transgene by qRT-PCR. We selected two overexpression lines (OE-*BnC07MYB3a-1*, OE-*BnC07MYB3a-2*) for subsequent analysis (Supplemental Fig. S3). Purple-black pigmentation was observed in the hypocotyls of 9-day-old black-seeded control material (cv. ZS11), but not in the OE-*BnC07MYB3a* lines (Fig. S4A). After DMACA staining, the hypocotyls of the OE-*BnC07MYB3a* lines were lighter than those of ZS11 (Supplemental Fig. S4A). Quantitative analysis further confirmed that the PA content in OE-*BnC07MYB3a* hypocotyls was significantly lower than that in ZS11 hypocotyls (Supplemental Fig. S4B).

The seeds of *OE-BnC07MYB3a* lines were lighter than the dark mature seeds of ZS11 (Fig. 4a). Near-infrared reflectance spectroscopy showed that OE-*BnC07MYB3a* seeds had higher degrees of the yellow-seeded trait than ZS11 seeds (Fig. 4B). DMACA staining revealed dynamic changes in PA contents in ZS11 and the transgenic lines during seed development. Both ZS11 and OE-*BnC07MYB3a* seeds were stained progressively darker blue as they developed, but the OE-*BnC07MYB3a* lines exhibited a lower degree of blue coloration than ZS11 seeds (Fig. 4C). The PA contents in the seed coats were consistent with the results of DMACA staining (Fig. 4, D-F).

**Figure 4.**
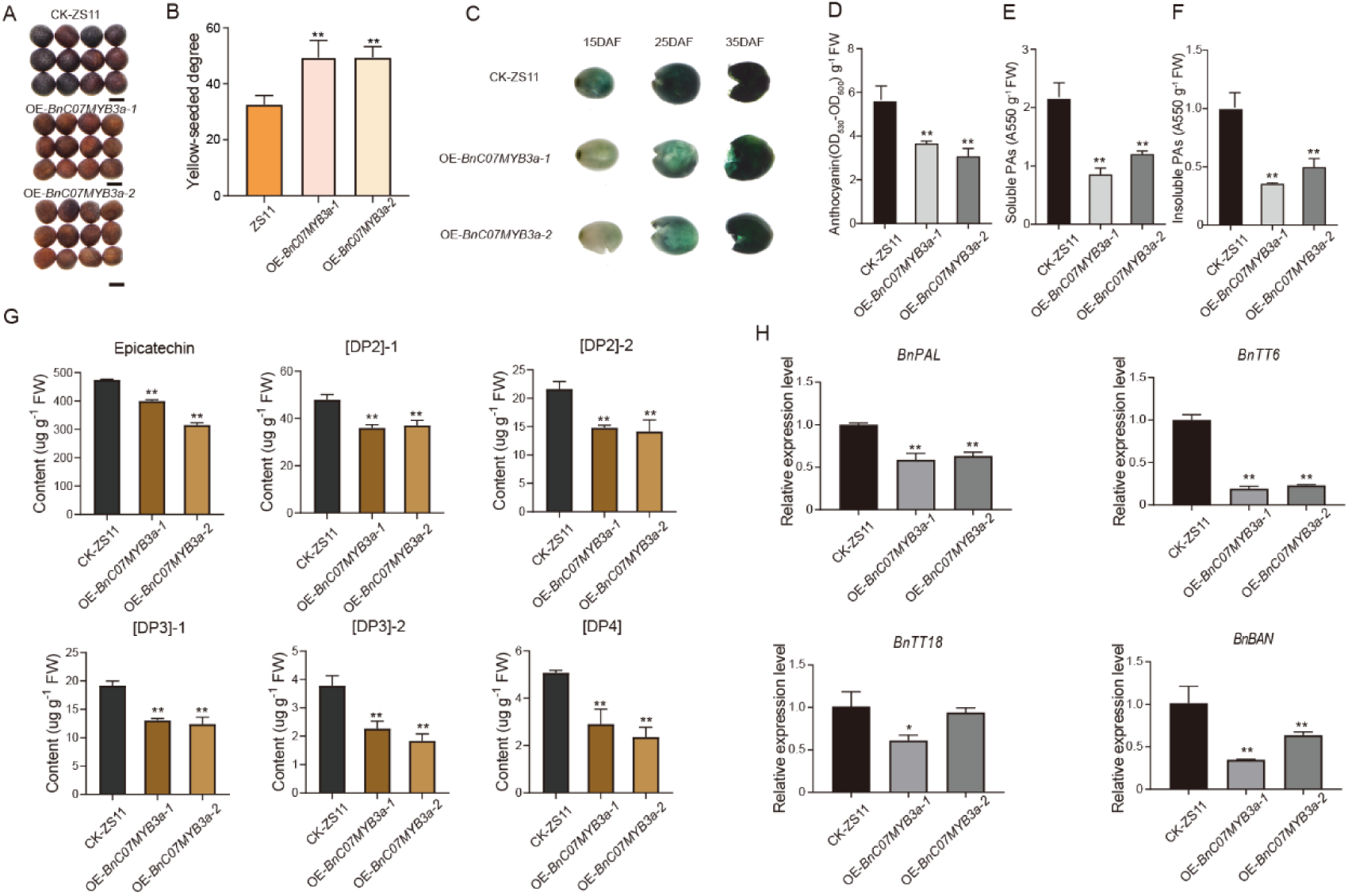
BnC07MYB3a negatively regulates the flavonoid biosynthesis pathway in rapeseed. A. Mature ZS11 and OE-*BnC07MYB3a* seeds. Bars, 1 mm. B. The yellow-seeded degree (YSD) of ZS11 and OE-*BnC07MYB3a* lines. (**, *p* < 0.01). C. DMACA staining of the seed coats of ZS11 and OE-*BnC07MYB3a* lines at different developmental stages. DAF, days after flowering. D. Anthocyanin levels in ZS11 and two transgenic lines. E. Soluble PA levels in ZS11 and two transgenic lines. F. Insoluble PA levels in ZS11 and two transgenic lines. G. Differentially abundant flavonoids identified by UPLC-HESI-MS/MS in ZS11 and OE-*BnC07MYB3a* seeds. H. Relative expression levels of flavonoid biosynthesis genes in ZS11 and OE-*BnC07MYB3a B. napus* lines, as detected by qRT-PCR. Values are mean ± SD of three biological replicates (**, *p* < 0.01; *, *p* <0.05). *P* values were calculated using multiple *t* tests without adjustments.

We quantified the flavonoid components in the transgenic lines by ultra-high-pressure liquid chromatography with heated electrospray ionization tandem mass spectrometry (UPLC-HESI-MS/MS). We detected significantly lower contents of epicatechin and PAs in the OE-*BnC07MYB3a* lines compared with ZS11 (Fig. 4G; Supplemental Fig. S5). qRT-PCR analysis revealed that *BnPAL*, *BnTT6*, *BnTT18*, and *BnBAN* expression was inhibited in the OE-*BnC07MYB3a* lines compared with ZS11 (Fig. 4H).

### BnC07MYB3a negatively regulates the expression of the flavonoid biosynthesis pathway gene *BnTT6*

Transcription factors inhibit or activate gene expression by binding to sequences upstream of the gene (Liu et al. 1999). We carried out DAP-seq assays to search for binding targets of BnC07MYB3a genome wide. The peak was concentrated on target gene transcriptional start sites (TSSs) (Fig. 5A). KEGG enrichment analysis showed that the BnC07MYB3a target sites were enriched for “Anthocyanin biosynthesis,” “Flavone and flavonol biosynthesis,” and “Flavonoid biosynthesis” (Fig. 5B). To identify the motifs of the BnC07MYB3a binding targets, were performed motif enrichment analysis using MEME-ChIP software. The most significantly enriched motif sequence was “AYCTAMCTAAYYAYM” (E-value=9.5×e^−1655^) (Fig. 5C), which was matched multiple times in the promoter region of *BnTT6* (Supplemental Fig. S6). Importantly, in an electrophoretic mobility shift assay (EMSA), recombinant glutathione S-transferase (GST)-BnC07MYB3 protein bound to a DNA probe containing the core motif (5′-AYCTAMCTAAYYAYM-3′). This signal decreased with increasing amounts of competitor probe, but not mutated probe, in the reaction (Fig. 5, D and E). In a dual-luciferase assay, co-transformation of the *BnTT6* promoter with BnC07MYB3a led to significantly less luciferase activity than co-transformation with the control, indicating that BnC07MYB3a regulates the expression of *BnTT6* (Fig. 5F). *BnTT6* was differentially expressed between two yellow-seeded lines (GH06 and L1188) and two black-seeded lines (ZY821 and ZS11), but we did not observe sequence differences in *BnTT6* between these different genetic backgrounds (Supplemental Fig. S7).

**Figure 5.**
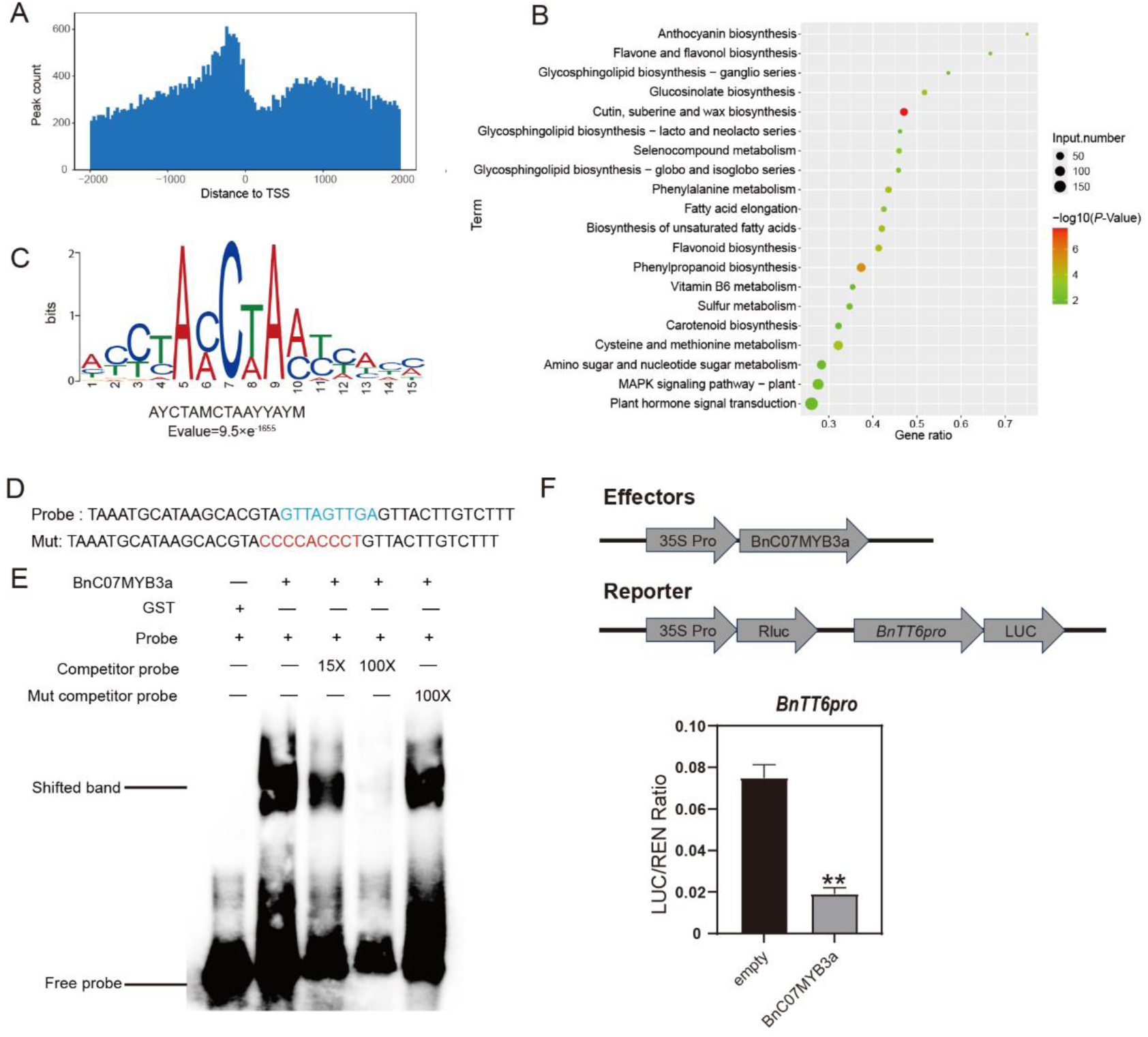
BnC07MYB3a directly regulates flavonoid pathway genes. A. BnC07MYB3a binding sites are centered on gene transcriptional start sites (TSSs) genome wide. B. KEGG enrichment analysis of putative BnC07MYB3a binding sites. C. The enriched DNA motif identified at BnC07MYB3a binding sites. E-value is shown. D. Probes used in the electrophoretic mobility shift assay (EMSA). The probe was labeled with biotin. E. EMSA of the binding of recombinant BnC07MYB3a protein to the promoter of *BnTT6*. F. Dual-luciferase reporter assay showing that BnC07MYB3a regulates *BnTT6* promoter activity. (**, *p* < 0.01). *P* values were calculated using multiple *t* tests without adjustments.

### BnA06bHLH92a and BnTTG1 interact with BnC07MYB3a to form the MBW complex

Numerous studies have shown that the MYB–bHLH–WDR (MBW) protein complex regulates the expression of structural genes in the flavonoid pathway and thus participates in pigment anabolism (Hichri et al. 2011; Xu et al. 2015; Colanero et al. 2018). We previously determined that BnA06bHLH92a, a key regulator of the proanthocyanidin pathway, interacts with BnTTG1 (Hu et al. 2023), and BnC07MYB3a was captured by BnA06bHLH92a during a yeast two-hybrid screening, suggesting that these three proteins might form a previously uncharacterized MBW complex in *B. napus*. Therefore, we examined the proteins that interact with BnC07MYB3a using a yeast two-hybrid assay, finding that BnC07MYB3a interacted with BnA06bHLH92a and BnA06TTG1a/BnC07TTG1b in yeast cells (Fig. 6A). In a bimolecular fluorescence complementation assay, strong yellow fluorescent signals were detected in the nucleus of *Nicotiana benthamiana* cells co-transfected with *A. tumefaciens* containing either nYFP–BnA06bHLH92a and cYFP–BnC07MYB3 or cYFP–BnC07MYB3 and nYFP–BnA06TTG1a/BnC07TTG1b (Fig. 6B). In addition, in a yeast three-hybrid assay, neither the induction of maltose binding protein (MBP) nor BnC07MYB3a protein expression in yeast cells affected the interaction between BnC08TT2 and BnTTG1, whereas the control MBP protein did not interact with BnC08TT2 or BnTTG1 (Fig. 6C).

**Figure 6.**
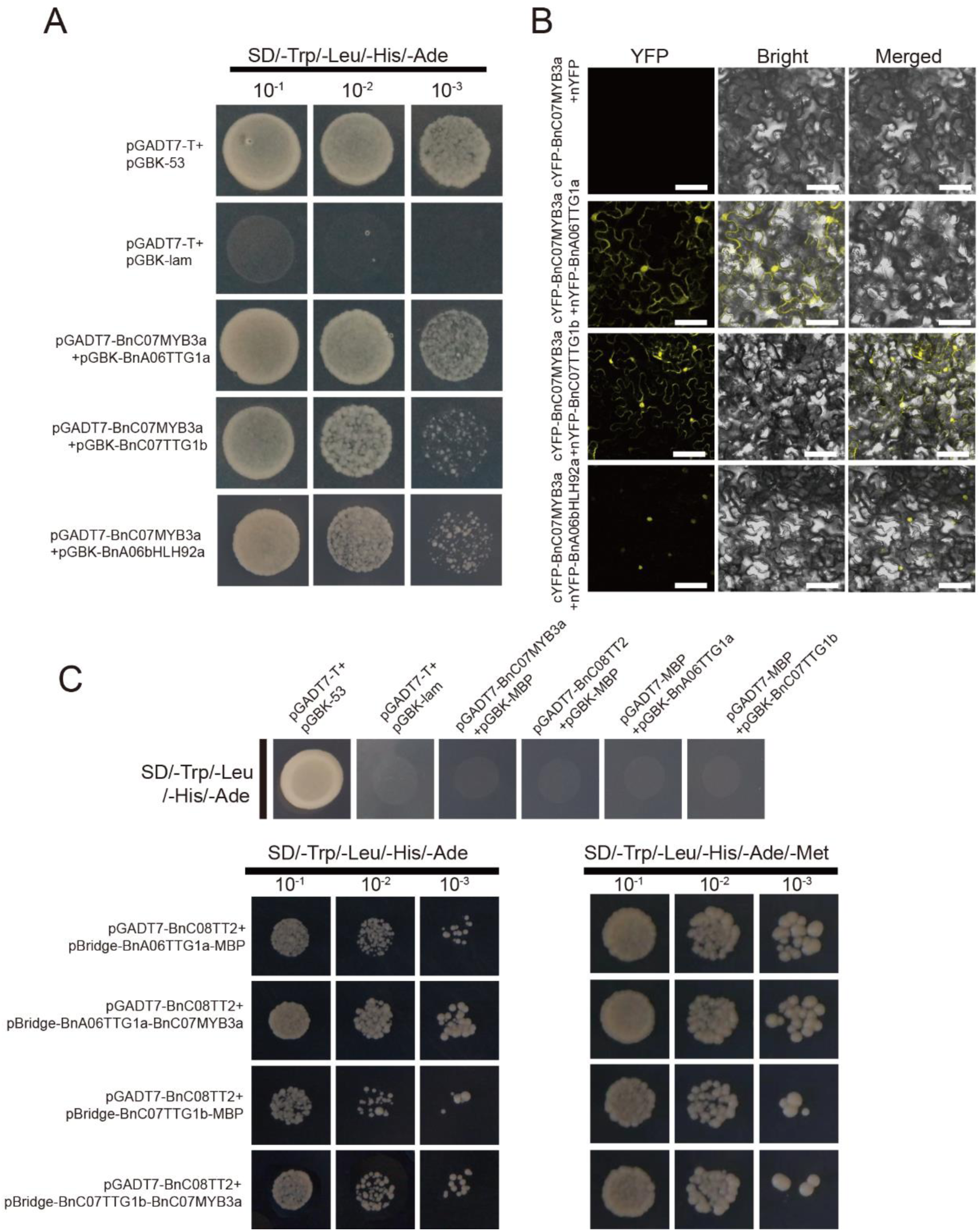
BnC07MYB3a interacts with BnTTG1 (BnA06TTG1a and BnC07TTG1b) and BnA06bHLH92a. A. Yeast two-hybrid assay of the interaction between BnC07MYB3a and BnTTG1/BnA06bHLH92a. pGADT7-T and pGBK-53, and pGADT7-T and pGBK-lam, were used as positive and negative controls, respectively. Ten-fold serial dilutions of transformed yeast cells were grown on SD-Trp/-Leu/-His/-Ade medium. B. Bimolecular fluorescence complementation assay of the interaction between BnC07MYB3a and BnTTG1/BnA06bHLH92a in *N. benthamiana* leaf epidermal cells. Bars, 50 μm. C. Yeast two-hybrid assay of the interaction between MBP with BnC07MYB3a, BnC08TT2 and BnTTG1. D. Yeast three-hybrid assays to verify whether BnC07MYB3a affects BnC08TT2 and BnTTG1 interactions. Yeast cells co-transformed with pGADT7-BnC08TT2 and pBridge-BnTTG1-BnC07MYB3a were grown on SD/-Trp/-Leu/-His/-Ade medium to assess BnC08TT2 and BnTTG1 interaction. Yeast cells co-transformed with pGADT7-BnC08TT2 and pBridge-BnTTG1-BnC07MYB3a were grown on SD/-Trp/-Leu/-His/-Ade/-Met medium to induce BnC07MYB3a expression and test for its effect on BnC08TT2 and BnTTG1 interactions. pGADT7-BnC08TT2 and pBridge-BnTTG1-MBP were used as controls.

## Discussion

The yellow-seed trait is particularly desirable in *B. napus*, as it is associated with higher oil and protein yields and improved seed quality (Meng et al. 1998; Tang et al. 1997). The yellow-seeded trait results from the absence or low accumulation of pigments in the seed coat, which is translucent and reveals the yellow color of the embryo (Zhai et al. 2020). Various flavonoids have been identified in yellow- and black-seeded *B. napus* using heated electrospray ionization tandem mass spectrometry (UPLC-HESI-MS/MS), indicating that the flavonoid pathway and its metabolites are the key factors affecting seed coat color (Jiang et al. 2013; Shao et al. 2014; Wang et al. 2018). Among these, epicatechin and its derivatives PAs are considered to be the main substances for the formation of seed coat color (Qu et al. 2013; Qu et al. 2020; Shen et al. 2021). Here, we detected great differences in the accumulation of epicatechin and its derivatives in six yellow- vs. black-seeded rapeseed materials (Fig. 1A). Transcriptome analysis of the *B. napus* materials revealed that the DEGs were enriched in the flavonoid pathway, which produces epicatechin (Fig. 1, B and C). These results underscore the significant role of epicatechin and its derivatives in seed coat pigment formation.

WGCNA is an effective gene mining method that is widely used in biological research (Langfelder et al. 2008). Here, we performed WGCNA using epicatechin and its derivatives as traits and analyzed the DEGs. We identified a red gene module containing 163 genes associated with the accumulation of epicatechin and its PA derivatives (Fig. 1, A–E; Supplemental Fig. S1). Using the STRING database to predict protein interactions, we identified a flavonoid network among these 163 genes, which included four transcription factor genes: *TT8*, *TT2*, *MYB3*, and *MYB63* (Fig. 1F). *BnTT8* and *BnTT2* influence the formation of seed coat pigments in rapeseed (Li et al. 2024), whereas *MYB63* and *MYB3* are involved in the lignin biosynthetic pathway, and *MYB3* also regulates anthocyanin accumulation in Arabidopsis (Zhou et al. 2009; Kim et al. 2022). These results indicate that the genes in the flavonoid network were accurately identified.

In the flavonoid biosynthetic pathway, MYB proteins can form a transcriptional acrtivation complex with bHLH and TTG1 (Xu et al. 2015). In this study, we determined that BnC07MYB3a contains a highly conserved R2R3 domain at its N-terminus and is highly homologous to AtMYB3, which negatively regulates anthocyanin/PA accumulation, suggesting that these transcription factors share a similar role (Supplemental Fig. S2). We determined that *BnC07MYB3a* was highly expressed in yellow-seeded varieties and encodes a transcriptional repressor, suggesting that BnC07MYB3a regulates seed coat color in *B. napus* (Fig. 2).

In a previous report, the expression levels of *DFR* and *LDOX* in Arabidopsis mutants *myb3* were examined using qRT-PCR, which did not provide a phenotypic picture of the pigmentation changes (Kim et al. 2022). To investigate the biological function of BnC07MYB3a, we generated transgenic Arabidopsis and *B. napus* lines that overexpressed *BnC07MYB3a*. Both OE-*3a* and OE-*BnC07MYB3a* showed lower anthocyanin and proanthocyanidin contents during seed development compared with the wild type, resulting in lighter seed coat color at maturity (Fig. 3; Fig. 4, A–F). UPLC-HESI-MS/MS analysis revealed that the levels of PA metabolites (epicatechin, [DP2]-1, [DP2]-2, [DP3]-1, [DP3]-2, and [DP4]) were significantly lower in the OE-*BnC07MYB3a* lines than in ZS11 (Fig. 4G; Supplemental Fig. S5), which is consistent with the previous finding that PAs are major constituents of dark seed coats in *Brassica* species (Qu et al. 2013; Shen et al. 2021). Flavonoid biosynthesis genes were also significantly repressed in seed coats of the OE-*BnC07MYB3a* lines (Fig. 4H), where their function in rapeseed was initially characterized (Li et al. 2024).

Direct binding of an R2R3-MYB repressor to the promoters of structural genes has rarely been demonstrated (LaFountain et al. 2021). We determined that BnC07MYB3a binds directly to the *BnTT6* promoter and inhibits its activity (Fig. 5; Supplemental Fig. S6). *BnTT6* encodes flavanone 3-hydroxylase, a precursor of flavonols and anthocyanins (Wang et al. 2021, Chen et al. 2024). TT6 is a key enzyme regulating flavonoid accumulation (Qu et al. 2013; Liu et al. 2022). Expression levels of *BnTT6* were significantly lower in the overexpression lines compared with those in ZS11, which could explain why the anthocyanin/proanthocyanidin content was reduced in these lines. Additionally, *BnTT6* was differentially expressed between yellow-and black-seeded *B. napus* accessions but did not display sequence differences (Supplemental Fig. S7), indicating that its regulation by BnC07MYB3a is responsible for the differences in seed coat pigmentation.

Flavonoid biosynthesis is regulated by activator-type MBW ternary complexes (R2R3-MYB, bHLH, and WD40), particularly the TT2-TT8-TTG1 complex (Lepiniec et al. 2006; Hichri et al. 2011). We determined that BnC07MYB3a physically interacts with BnTTG1 (BnA06TTG1a and BnC07TTG1b) and BnA06bHLH92a (Fig. 6, A and B), a negative regulator of anthocyanin and proanthocyanidin accumulation (Hu et al. 2023). These findings suggest that BnC07MYB3a, BnA06bHLH92a, and BnTTG1 form a previously uncharacterized MBW complex. Several studies have shown that repressive MYB transcription factors interact with core MBW complex members, which interferes with the regulation of the flavonoid pathway by the MWB complex (Xie et al. 2022; Xiang et al. 2019). By contrast, BnC07MYB3 does not affect MBW complex formation by interacting with BnTTG1 (Fig. 6C).

In this study, we revealed the role of BnC07MYB3a as a negative regulator of the anthocyanin and PA biosynthesis pathways. In addition, we provided molecular evidence that BnC07MYB3a inhibits the expression of *BnTT6*, thus leading to decreased flavonoid metabolite accumulation and a lighter-color seed coat. Based on these findings, we propose a model for the role of BnC07MYB3a in regulating the anthocyanin and PA biosynthetic pathways to determine seed coat color (Fig. 7). Our results lay the foundation for elucidating the regulatory mechanisms of anthocyanin and PA biosynthesis in *B. napus* and other important crops.

**Figure 7.**
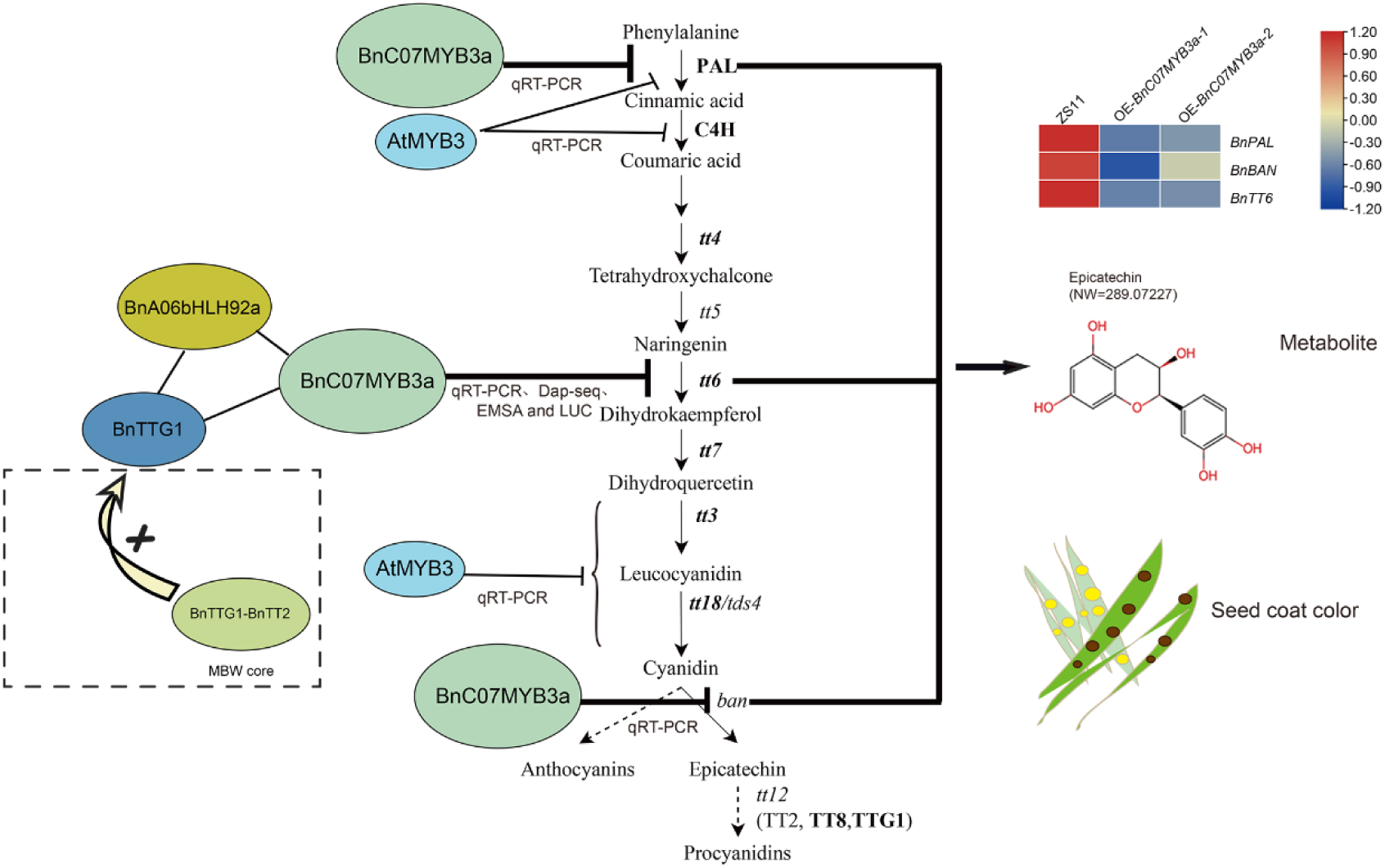
BnC07MYB3a represses the expression of the flavonoid pathway gene *BnTT6* to reduce anthocyanin and PA accumulation. Based on qRT-PCR in OE-*BnC07MYB3a* lines, *BnBAN* and *BnPAL* may be target genes of BnC07MYB3a. Based on qRT-PCR in Arabidopsis mutants *myb3* lines, *PAL, C4H, TT3 (DFR)* and *TT18 (LDOX)* may be target genes of AtMYB3. BnC08TT2 and BnTTG1 are core components of the MBW complex, which affects anthocyanin and PA accumulation by regulating the expression of late biosynthetic genes in the flavonoid pathway.

## Materials and methods

### Plant materials and growth conditions

The *Brassica napus* materials used in this study were the black-seeded cultivar Zhongshuang11 (ZS11) and ZY821, the yellow-seeded accession GH06 and L1188, and the transgenic line OE-*BnC07MYB3a*, which were planted in the transgenic base of Southwest University in Chongqing, China. Five well-developed *B. napus* plants were randomly selected from each line. The flowering dates at the full bloom stage were marked using different colors of wool, and seeds from different developmental stages were collected from each line based on the flowering date. The seeds were quickly frozen in liquid nitrogen and stored at −80℃.

*Arabidopsis thaliana* ecotype Columbia-0 (Col-0) and *Nicotiana benthamiana* were grown in a light incubator under a 16-h-light/8-h-dark cycle at a daytime temperature of 22°C for Arabidopsis and 25°C for *N*. *benthamiana* and a nighttime temperature of 20°C, with a relative humidity of 40%–65% and a light intensity of 16,000 lx.

### RNA-seq

Total RNA was extracted from the seeds of *B. napus* during different developmental stages using an EZ-10 DNAaway RNA Mini-Prep Kit (Sangon, Shanghai, China) following the manufacturer’s instructions. Transcriptome sequencing was performed on the Illumina HiSeq2000 sequencing platform at Novogene Bioinformatic Technology Co., Ltd. (Tianjin, China). Clean reads were obtained by removing low-quality reads containing adapters or poly-N from the raw data (Bolger et al. 2014) and mapped to the assembled high-quality genomes of yellow-seeded *B. napus* GH06 and black-seeded *B. napus* ZY821 (https://www.ncbi.nlm.nih.gov/bioproject/PRJNA770894) (Qu et al. 2023). Gene expression levels were calculated based on FPKM (fragments per kilobase of transcript per million fragments mapped) values; differential gene expression analysis was performed using DEGseq (version 1.44.0) (https://bioconductor.org/packages/DESeq2/). The threshold of DEGs was fold change ≥ 2 and *p*-value ≤ 0.05.

### Co-expression network construction by WGCNA

Gene co-expression networks were constructed using the WGCNA package (version 1.72) with RNA-Seq data from seeds (Langfelder et al. 2008). DEGs or genes in each module were subjected to Kyoto Encyclopedia of Genes and Genomes (KEGG) pathway analysis using the R package clusterProfiler (Yu et al. 2012). The correlation between eigengenes and metabolites in the modules was calculated using a Pearson test. Subsequently, the protein sequences of the red module genes were submitted to STRING (https://string-db.org/cgi/input.pl) and plantTFDB (http://planttfdb.gao-lab.org/) (Tian et al. 2020) websites for Protein-Protein interactions network analysis and transcription factor prediction, respectively. And Cytoscape ver. 3.5.1 (https://cytoscape.org/) was used to visualization of Protein-Protein interactions network.

### Gene cloning, phylogenetic analysis, and expression analysis

Total RNA was extracted from *B. napus* plants using an EZ-10 DNAaway RNA Mini-Prep Kit (Sangon Biotech, Shanghai, China) following the manufacturer’s instructions. Full-length *BnMYB3a* cDNA was amplified from ZS11 cDNA using TransStart FastPfu Fly DNA Polymerase (TransGen, Beijing, China). Multiple sequence alignment was performed using COBALT (Constraint-based Multiple Alignment Tool; https://www.ncbi.nlm.nih.gov/), and a phylogenetic tree was reconstructed using MEGA7 (Lynnon Biosoft, Quebec, Canada) with the neighbor-joining method (Kumar et al. 2016). Quantitative reverse-transcription PCR was conducted in a CFX96 real-time PCR machine (Bio-Rad Laboratories, Hercules, CA, USA) as described previously (Hu et al. 2023). Housekeeping genes (*BnActin7*, *AtACTIN2*) were used as internal reference genes during data analysis. The primers used in the experiment are listed in Table S2.

### Transformation of Arabidopsis and *B. napus* with *BnC07MYB3a*

The coding sequence of *BnC07MYB3a* was cloned into the pNC-Cam3304-MCS35S vector driven by Pro35S (Yan et al. 2019) and into the pCAMBIA-1303 vector driven by P-GLY (Zhang et al. 2020b). The above recombinant plasmids were transformed separately into *Agrobacterium tumefaciens* strain GV3101. Agrobacterium containing recombinant plasmids was infiltrated into Arabidopsis (Col-0) flowers and *B. napus* hypocotyl tissues to obtain overexpression plants as described previously (Lu et al. 2013, Qi et al. 2019). After spraying the leaves of transgenic plants with 100 mg L^-1^ Basta, the presence of the transgene was confirmed by qRT-PCR. Plants with normal growth and development were selected, and transgene-positive homozygous progenies were used for subsequent analyses. The primers used in the experiment are listed in Table S2.

### Subcellular localization of BnC07MYB3a

The coding region of *BnC07MYB3a* was amplified by PCR and cloned into pNC-Cam1304-SubN (Yan et al. 2019), containing a *GFP* gene at the N-terminus, to generate the Pro35S-BnC07MYB3a::GFP fusion vector. The recombinant plasmid was transformed into Arabidopsis protoplasts. pNC-Cam1304-SubC (GFP) was used as a positive control. Fluorescent signals were observed under a confocal laser-scanning microscope (LSM802400301, Carl Zeiss, Oberkochen, Germany) 2 days after infiltration. The primers used in the experiment are listed in Table S2.

### Yeast two-hybrid and yeast three-hybrid assays

To identify proteins that interact with BnC07MYB3a, yeast two-hybrid (Y2H) assays were performed using the Matchmaker Gold Y2H System (Clontech, USA). The full-length coding sequences of *BnC07MYB3a*, *BnC08TT2*, and maltodextrin binding protein (*MBP*) were each ligated into pGADT7. The full-length coding sequences of *BnA06bHLH92a*, *BnTTG1* (*BnA06TTG1a*/*BnC07TTG1b*), and *MBP* were each ligated into pGBKT7. The AD and BD fusion vectors were introduced into yeast (*Saccharomyces cerevisiae*) strain Y2H Gold using the polyethylene glycol/lithium acetate method described in the Yeast Protocols Handbook (Clontech). Co-transformed colonies were initially grown separately on SD/-Trp/-Leu plates and further screened for growth on SD/-Trp/-Leu/-His/-Ade plates. The pGADT7-T and pGBKT7-53 vectors were used as a positive control. The pGADT7-T and pGBK-lam vectors were used as a negative controls. All experiments were repeated three times.

To investigate whether BnC07MYB3a interferes with the MBW complex, yeast three-hybrid (Y3H) assays were carried out based on the Y2H experimental system. The coding region of *BnA06TTG1a* or *BnC07TTG1b* was recombined into multiple cloning site I (MCS I) of the pBridge vector, and the coding region of *BnC07MYB3a* was recombined into multiple cloning site II (MCS II) of the pBridge vector, resulting in the pBridge-BnA06TTG1a-BnMYB3a or pBridge-BnC07TTG1b-BnMYB3a fusion vector. MBP, which does not interact with BnC08TT2 or BnTTG1, was used as a negative control. The interactions among fusion proteins expressed from the recombinant pBridge and pGADT7 vectors were analyzed as described previously (Chakravorty et al. 2015). The primers used in the experiment are listed in Table S2.

### Bimolecular fluorescence complementation (BiFC) assay

The full-length coding sequences of *BnC07MYB3a* and *BnA06bHLH92a* were cloned into pNC-BiFC-Ecc (Yan et al. 2019), containing the C-terminal fragment of YFP (cYFP) or pNC-BiFC-Enn (Yan et al. 2019), containing the N-terminal fragment of YFP (nYFP), respectively, resulting in the recombinant plasmids cYFP-BnC07MYB3a and nYFP-BnA06bHLH92a. The recombinant plasmids nYFP-BnA06TTG1a and nYFP-BnC07TTG1b were constructed as described previously (Hu et al. 2023). *Agrobacterium tumefaciens* cells containing recombinant plasmids were co-injected into the leaves of 6-week-old *N. benthamiana* plants in different combinations. Fluorescent signals were observed under a confocal laser-scanning microscope (LSM802400301, Carl Zeiss, Oberkochen, Germany) 2 days after infiltration. The primers used in the experiment are listed in Table S2.

### Transient dual-luciferase expression assays

To investigate the transcriptional repressor activity of BnC07MYB3a, the VP16 vector harboring the full-length cDNA of *BnC07MYB3a* and empty VP16 vector were used as effectors. The reporter vector contained the GAL4 binding element and the minimal CaMV 35S promoter fused to the firefly luciferase (*LUC*) reporter gene, whereas the Renilla luciferase (*REN*) reporter gene under the control of the 35S promoter was used as an internal control. The promoter sequence of *BnTT6* was cloned into the pNC-Green LUC vector (Yan et al. 2019) to generate BnTT6pro-LUC as a reporter. The coding region of *BnC07MYB3a* was cloned into the pNC-Green-SK vector (Yan et al. 2019) to generate the effector construct SK-BnC07MYB3a. Arabidopsis protoplasts were transformed with the effector and reporter constructs using an Arabidopsis Protoplast Preparation and Transformation Kit (Coolaber). Luciferase activity was determined using the Dual-Glo Luciferase Assay System (Promega) in a luminescence detector (Promega, GloMax 20/20) after 12–16 h. At least six biological replicates were conducted for each combination. The primers used in the experiment are listed in Table S2.

### Seed colour measurement, DMACA staining, measuring PA content, and UPLC−HESI−MS/MS analysis

The yellow-seeded degrees (YSD) was calculated using the near-infrared reflectance spectroscopy (DS2500, Foss Analytical A/S) and used to evaluate the seed colour phenotype according to the method as described previously (Fu et al. 2007; Daszykowski et al. 2008). Embryos of the control material ZS11 and the overexpression lines were removed from seeds at different developmental stages, and the seed coats were retained for DMACA staining. Staining was carried out with 5% (w/v) 4-dimethylaminocinnamaldehyde (DMACA) solution as described previously (Hong et al. 2017). Digital images were acquired under an Olympus SZX7 stereomicroscope (Olympus, Tokyo, Japan) and processed using Adobe Photoshop CS6.0 software (Adobe, San Jose, USA). Soluble and insoluble proanthocyanidins were extracted from *B. napus* seeds using the butanol-HCl method, and absorbance values were measured at a wavelength of 550 nm using a spectrophotometer (Liang et a., 2006). Proanthocyanidin levels (soluble and insoluble) are represented by A_550_ (g FW)^-1^.

Fresh seeds (100 mg) were quickly crushed into a powder and dissolved in an aqueous solution of formic acid (0.1%, v/v) in aqueous methanol (1 mL; 80%, v/v). The samples were shaken in an ultrasonic cleaner at room temperature for 1 h. The supernatant was filtered through a 0.22-μm nylon filter and subjected to UPLC-HESI-MS/MS analysis as described previously (Qu et al. 2020). The raw MS data have been deposited in MetaboLights under accession number MTBLS6703 (https://www.ebi.ac.uk/metabolights/search). Chemical structures were retrieved from the PubChem database (https://pubchem.ncbi.nlm.nih.gov). At least three biological replications were performed.

### DNA-affinity purification sequencing

DAP-seq binding assays were performed as described previously (Bartlett et al. 2017, Tang et al. 2021). Genomic DNA was extracted from *B. napus* leaves using the CTAB method, and an appropriate amount of genomic DNA was collected for fragmentation and screening. Subsequently, an affinity purification library was constructed from the screened genomic DNA using an NGS0602-MICH TLX DNA-Seq Kit according to the manufacturer’s instructions (Mich Scientific, Hebei, China).

Full-length *BnC07MYB3a* cDNA was cloned into the pFN19K HaloTag T7 SP6 Flexi expression vector. In vitro expression of Halo-BnC07MYB3a fusion proteins was performed using the TNT SP6 Coupled Wheat Germ Extract System (Promega, USA) according to the manufacturer’s instructions. The fusion proteins were directly purified using Magne Halo Tag Beads (Promega, USA). HaloTag–transcription factor fusion proteins were incubated with an adaptor-ligated genomic DNA library, and unbound DNA fragments were washed away. The samples were heated to release transcription-factor-bound DNA complexes, the supernatant was collected for next-generation sequencing, and the resulting genome-wide binding events were analyzed. And the DAP-seq datasets were supplied to MACS2 for peak calling. Association of peaks located 2 kb upstream or downstream of the transcription start sites (TSSs) were analyzed using Homer. Motif discovery was performed using the MEME-ChIP suite 5.0.5.

### Electrophoretic mobility shift assay (EMSA)

To produce glutathione S-transferase (GST)-tagged proteins, *BnC07MYB3a* was amplified, cloned into the pGEX4T-1 vector, and expressed in *Escherichia coli* strain Rosetta. Recombinant GST-BnC07MYB3a protein was purified using GST-tag Purification Resin (Beyotime, Jiangsu, China). EMSA was performed using a LightShift Chemiluminescent EMSA kit (Thermo Fisher Scientific, Waltham, MA, USA). The primers used in the experiment are listed in Table S2.

## Author contributions

R.H., C.Q. and J.L. conceived and designed the entire research program. L.G. and M.Z. culture transgenic materials. R.H., M.Z., M.T. and Y.L. completed most of the experiments. M.Z. and Y.L. watering plant material. S.S. and H.L. performed bioinformatics analysis. L.L., H.W. and K.L. provided to germplasm resources. R.H., and S.S. wrote original draft. R.H., C.Q., J.L., H.D., N.Y. and H.Z. performed editing and review. All authors read and approved the final article.

## Supplemental data

Additional supporting information can be found online in the Supplemental data section at the end of this article.

**Figure S1.** Cluster dendrogram and module colors of 1942 DEGs.

**Figure S2.** Characterization of BnC07MYB3a proteins.

**Figure S3.** Relative expression levels of BnC07MYB3a in OE-*BnC07MYB3a* rapeseed lines.

**Figure S4.** BnC07MYB3a inhibits PAs accumulation on the hypocotyl of *B. napus* seedlings.

**Figure S5.** UPLC-HESI-MS/MS chromatograms of significant constituents detected in the seeds of control and transgenic plants.

**Figure S6.** BnC07MYB3a binding motifs in the *BnTT6* promoter.

**Figure S7.** Phenotypes and characterization of *BnTT6*.

**Table S1**. Sequences information of *BnC07MYB3a*, *BnA06bHLH92a*, *BnTTG1*(*BnA06TTG1a and BnC07TTG1b*), *BnC08TT2* and *BnTT6* promoter in this study.

**Table S2.** Primers used in this study.

